# Locomotion Efficiency of Elephants: Mechanical work and energetics

**DOI:** 10.1101/2024.05.03.592175

**Authors:** Joakim G. Genin, Arthur H. Dewolf, Patrick A. Willems, Norman C Heglund

**Affiliations:** Laboratory of biomechanics and Physiology of Locomotion, Institute of NeuroScience, Université catholique de Louvain, Louvain-la-Neuve, Belgium

**Keywords:** elephant, locomotion, mechanical work, muscular efficiency

## Abstract

The size scaling of energy expenditure during locomotion has long puzzled researchers seeking biological invariants. Existing data, most of it from animals smaller than 200 kg, show that the mass specific mechanical work of locomotion is nearly independent of size, yet the metabolic cost of locomotion decreases with increasing size. Major questions remain concerning heavier animals: extrapolating mechanical work and metabolic cost to animals the size of an elephant comes to the uncomfortable conclusion that elephants would produce more energy than they would consume. Our study addresses this longstanding size-scaling conundrum by focusing on the locomotion mechanics and energetics of elephants, the largest extant land animals. In this study, the work required to move the limbs relative to the centre of mass of the whole body (*COM*) was measured in 27 Asian elephants (872-4000 kg). The total mechanical work was calculated by adding the external work required to maintain the movements the *COM*. Our investigation challenges the belief that mass-specific mechanical work of locomotion is independent of size. Furthermore, our study unveils a surprising aspect of elephant locomotion—the substantial fraction of total mechanical work dedicated to limb swinging. At high speeds, elephants allocate ∼75% of their work to limb swinging. Also, by quantifying the metabolic cost and mechanical work, we provide the first estimation of the efficiency of converting metabolic energy into mechanical work in elephants. Elephants exhibit an efficiency of at best 45%, which is similar to that in other large mammals such as horses and humans. This study not only addresses a significant gap in our understanding of size scaling in locomotion but also opens new avenues for exploring the evolution of energy expenditure across diverse animal sizes.

## Introduction

Terrestrial movement energetics and its relationship with size have captivated biologists for more than 50 years[1]. One major question is related to the quest of biological invariants, trying to explain and predict the evolution of the cost of locomotion. In 1982, Taylor et al. measured the net metabolic cost of transport (*E*_metab_, in J kg^--1^ m^-1^) of mammals and birds with a body mass range of 0.01 to 260 kg [2]. They found that the mass-specific *E*_metab_ was roughly inversely proportional to the cube root of the body mass (*M*_b_), *i*.*e. E*_metab_ ∝ *M*_b_^-0.32^. The same relationship was later found in reptiles, amphibians and arthropods weighing less than 10 g [3].

According to the above relation, since the elephant is the biggest living terrestrial animal, it should also be the most economical. Langman et al. [4,5] confirmed this hypothesis. They measured the energy expenditure during locomotion of African elephants weighing about 3.5 tons. These authors found that elephants present the lowest mass-specific metabolic cost of terrestrial locomotion ever measured: the addition of animals with a 12-fold increase in body mass over the largest animal measured by Taylor et al. [2] produced an allometric relationship for mammals ranging from 7g to 3.5 tons (*E*_metab_ ∝ *M*_b_^-0.27^). In other words, the mass-specific cost per unit distance of an elephant is ∼1/40 of the cost of a mouse.

It is commonly accepted that the main factor determining the energy expenditure of terrestrial locomotion is the mechanical work done by the animal to travel along the land [6,7]. During legged locomotion, the limbs (i) support and propel the body by exerting forces against the ground and (ii) oscillate forwards to be ‘reset’ each step. For this reason, the muscular work of locomotion falls naturally into two parts: (i) the work required to maintain the movements of the centre of mass (*COM*) relative to the surroundings (defined as external work, *W*_ext_) and (ii) the work required to maintain the movements the body segments relative to the *COM* (defined as internal work, *W*_int_). The total mechanical work (*W*_tot_) done during locomotion is then best computed as the sum of *W*_ext_ and *W*_int_ [8,9].

In the early eighties, Heglund and his colleagues publishes a series of four papers, in which they measured *W*_int_, *W*_ext_ and *W*_tot_ in different species of birds and mammals with a body mass ranging from 0.01 to 260 kg [2,10–12]. They found that the mass specific mechanical work that muscles perform to swing the limbs and raise and lower the *COM* does not vary with size. Extrapolating the metabolic and mechanic cost of locomotion to an animal the size of an elephant would come to the unrealistic conclusion that a 1500 kg or larger elephant would be producing more energy than it consumed.

Two papers evaluated the mechanical work in elephants [13,14], considering the work required to move the COM whereas the second one considered an individual limb method (inverse dynamics). However, neither of them considered the cost of swinging the limb (*W*_int_). Although *W*_int_ represents a significant fraction of *W*_tot_ during steady state level locomotion, particularly at high speeds [15], its impact on the total energy demand during elephant’s locomotion is still unknown. Taylor et al. [16] found that, despite large differences in limb mass, the energetic cost of running in cheetahs, gazelles, and goats of about the same total body mass and limb length was nearly identical over a wide range of speeds, suggesting that most of the energy expended in running might not be used to accelerate and decelerate the limbs. Furthermore, in their attempt to find a unifying hypothesis to explain simply the cost of locomotion, Kram and Taylor assumed that the cost of swinging the limbs back and forth was negligible [17]. More recently, however, Marsh et al. [18] found that the metabolic cost of swinging the limbs in guinea fowl was far from negligible, comprising 25% of total energy expenditure, while the cost of generating force to support the body during the stance phase accounts for 75% of the total.

Compared with smaller animals, elephants have heavy and bulky legs, which together increase inertia of the limbs. Thus, the elephant is an extreme case which may highlight the importance of *W*_int_ on *W*_tot_ during terrestrial locomotion.

Hildebrand and Hurley (1985) studied elephants at near their maximal speed [19] and found that large oscillations in the kinetic energy of the limbs occur both during stance and swing phase of the limb. Unfortunately, a large amount of information is missing from this study (weight of the animals, the energy curves are not normalised, unknown running speed, no evolution of the energy at different speeds, no description of the determination of the position of the *COM* for each segment, unknown reference point for energy fluctuations, the head and torso segment are not considered, and so on), making comparisons to this study impossible.

Furthermore, elephants move with a high step frequency compared to other large animals at the same speed. Genin et al. [13] have observed that the highest speed of an elephant corresponds to a slow trot for a 2-ton quadruped, while the step frequency adopted is higher than the maximum predicted step frequency for an animal of that size, even at a top speed gallop. In humans, walking or running at a given speed with a higher step frequency requires less *W*_ext_ but more *W*_int_ [9,20–22]. Since at a given speed elephants are moving with higher step frequency than other large animals, the relative contribution of *W*_int_ to *W*_tot_ could also be modified.

In this study, the kinetic energy of the body segments relative to the *COM* in Asian elephant is measured. The internal work is calculated from these energy curves, and together with the measurements of the external work [13], a complete figure of the total mechanical work is presented and compared to the individual limb methods in [14]. Finally, the total muscular work measured in this study is compared with the metabolic cost measured by Langman et al. [4,5], to estimate the efficiency of positive work production during elephant locomotion. Work, cost and efficiency will be compared to allometric predictions, opening new directions for re-examining the comparative mechanical costs of terrestrial locomotion and revising classical scaling theories about locomotor costs and efficiency.

## Materials and Methods

The internal work, *W*_int_, was measured on 27 Asian elephants (*Elephas maximus*), at the Thai Elephant Conservation Centre, near Lampang in Northern Thailand. Experiments were approved by the Ethic Committee of the Faculty of Medicine of the UCL and by the Forest and Industry Organization (FIO) in Thailand.

The body mass (*M*_b_) of the elephants ranged from 1.3 to 4 tons. Morphological values for each elephant are listed in Table 1. In most cases, a mahout representing 1-5% of the weight of the elephant rode on the elephant; we assumed the mahout had no influence on movements of the elephant’s limbs.

**Table 1.** Individual data for elephants used in this study. Maximal forward velocity for each elephant and the corresponding Froude number are presented for comparison with the literature.

| Elephant | Sex | Age<br>(yrs) | Body<br>Mass<br>$M_b$ (kg) | Mass on the<br>front legs<br>(% $M_b$) | Hip Height<br>(m) | Maximal<br>velocity<br>(m s<sup>-1</sup>) | $Fr$ |
| --- | --- | --- | --- | --- | --- | --- | --- |
| Add | M | 8 | 2515 | 57.4 | 1.64 | 2.04 | 0.70 |
| Boonmii | F | 43 | 3376 |  | 1.79 | 2.8 | 1.43 |
| Gaew | M | 5 | 1740 | 59.4 | 1.52 | 4.97 | 3.83 |
| Janpui | M | 36 | 3259 |  | 1.72 | 2.52 | 1.11 |
| Jojo | M | 13 | 3000 | 59.3 | 1.77 | 4.51 | 3.67 |
| Kampaeng | M | 37 | 3221 |  | 1.66 | 2.25 | 0.86 |
| Kangluay | M | 9 | 1956 | 62.1 | 1.61 | 1.81 | 0.54 |
| Kumlha | M | 58 | 3125 | 60.2 | 1.84 | 1.31 | 0.32 |
| Lookkhang | F | 12 | 2635 | 62.2 | 1.72 | 4.61 | 3.73 |
| Monkol | M | 29 | 3147 |  | 1.8 | 1.11 | 0.23 |
| Phajan | M | 45 | 3869 | 61.2 | 1.83 | 2.06 | 0.79 |
| Pong | M | 7 | 2274 |  | 1.59 | 2.64 | 1.13 |
| Prajuab | F | 24 | 3061 | 58.8 | 1.7 | 2.42 | 1.01 |
| Prame | M | 43 | 3218 | 59.5 | 1.84 | 1.68 | 0.53 |
| Pratida | F | 11 | 2617 |  | 1.69 | 2.02 | 0.70 |
| Prayao | M | 24 | 4001 |  | 1.86 | 2.58 | 1.26 |
| Satit | M | 29 | 3525 |  | 1.89 | 1.98 | 0.76 |
| Somboon | M | 53 | 3131 | 59.7 | 1.7 | 1.34 | 0.31 |
| Srisiam | M | 6 | 1258 | 63.3 | 1.32 | 4.85 | 3.17 |
| Tadaeng | M | 36 | 3543 | 62.8 | 1.71 | 0.82 | 0.12 |
| Tantawan | F | 47 | 3636 |  | 1.7 | 1.29 | 0.29 |
| Tao | M | 7 | 1893 | 64.7 | 1.58 | 1.72 | 0.48 |
| Umpang | F | 8 | 1641 | 59.6 | 1.32 | 3.58 | 1.72 |
| Wanalee | F | 8 | 2072 | 63.2 | 1.51 | 4.32 | 2.87 |
| Wangjao | F | 45 | 3102 |  | 1.58 | 1.72 | 0.48 |
| Wassana | M | 48 | 3078 | 62.6 | 1.8 | 1.13 | 0.23 |
| Yai | M | 48 | 3034 | 58.9 | 1.72 | 1.23 | 0.27 |
| Mean |  | 27.4 | 2849 | 60.9 | 1.68 | 2.42 | 1.2 |
| SD |  | 17.7 | 712 | 2 | 0.15 | 1.24 | 1.2 |

Elephants moved at different speeds along a 50 m-long track. Measurements of *W*_int_ were realised simultaneously with the measurement of the external work, *W*_ext_. The external work was evaluated by means of a 2 x 8 m force-platform mounted at ground level in the middle of the track. The method to compute *W*_ext_ from the ground reaction forces and the results obtained on *W*_ext_, were presented in Genin et al. [13].

Before data acquisition, elephants walked a few times back and forth on the track to become familiar with the experimental set-up. At the beginning of the data acquisition, the elephant moved at their freely chosen speed. During the next trials, the mahout encouraged the elephant to go slightly faster, up to the maximal speed. At the highest speeds, elephants were motivated by playful chasing or by food rewards at the end of the track. The data acquisition ended with the trials at the slowest speeds. During all the trials, the average speed of the elephants was measured by means of two photocells placed at each end of the force-platform and adjusted at the level of the elephant’s forehead.

### Measurements of the limb movements

The movements of the limb segments of the elephants were recorded using a high-speed digital camera A501k Basler^®^ (Vision Technologies, Ahrensburg, Germany). The resolution of the camera was 1280 x 1024 pixels and the aperture time of the objective was 3 ms. According to the speed of progression, images were sampled at a rate of 50-100 frames s^-1^. The camera was positioned perpendicular to the middle of the 8 m long force-plate, 15.3 m to the side and 1.15 m above the plate surface. The field of the camera encompassed about 12 x 4 m and its centre pointed roughly at the centre of the torso of an elephant that would stand in the middle of the force plate.

The body of the elephant was divided into 14 rigid segments: two hindfeet, two shanks, two thighs, one torso, two upper foreleg, two lower forelegs, two forefeet and one head. The limb segments facing the camera were delimited by white dots painted at the level of the lateral surface of the joints (Fig. 1). The marker painted in the middle of the torso also allowed evaluating the movements of the hip and the shoulder relative to that marker.

**Figure 1:**
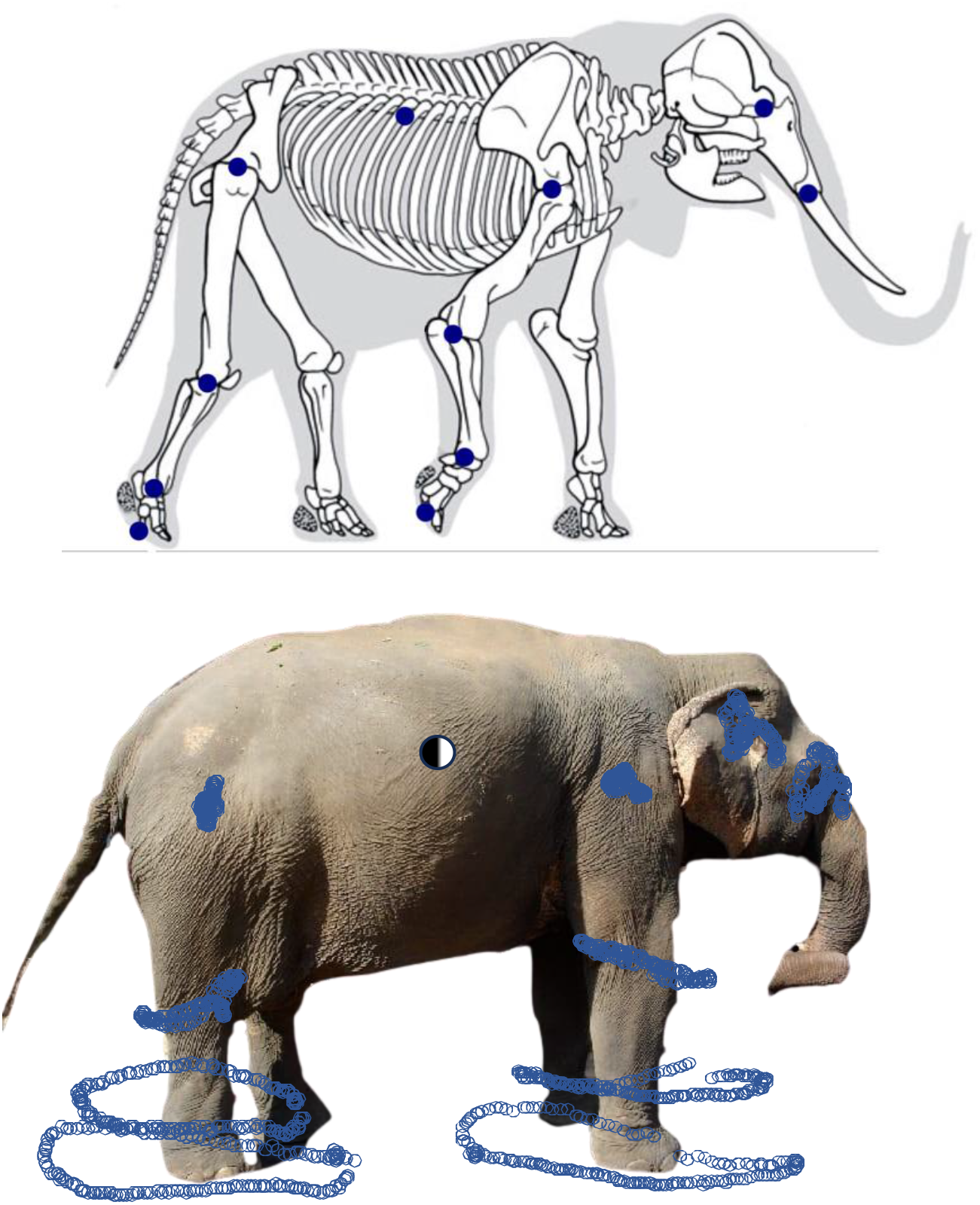
Marker placements on the right side of a representative elephant’s skeleton with skin contour in greyed background. These markers were used to determine the position of the segments. Joint centre positions were estimated by palpation and by movements of flexing-extending limbs. Picture from (Shoshani and Knight, 1992).

The coordinates of the markers in the fore-aft and vertical directions were measured each frame using Lynxzone software (Arsalis, Glabais, Belgium). Displacements in the lateral direction were ignored. The images were calibrated using the dimensions of the force-platform. The error due to parallax on the distances and angles measured at the centre of the image compared to that measured at the side of the image represents less than 3% of the measurement. In order to minimise this error, measurements were realised only on strides recorded close to the centre of the image. Strides were delimited by the right hind-limb touchdown. For each elephant, one stride per speed class of 0.28 m s^-1^ was analysed, throughout its entire range of speed.

### Calculation of the internal work Wi_nt_

The mechanical internal work (*W*_int_) done to move the body segments relative to the *COM* was evaluated from their translational and rotational kinetic energy, using a procedure similar to that described in detail by Willems et al. [8]. Only translational and rotational movements taking place in the sagittal plane were considered.

The angle made by each segment with the horizontal was calculated each frame from the coordinates of the markers. The angle-time curves were smoothed using a spline function (Matlab®, Mathworks, Natick, MA, USA). A ‘stick elephant’ was then reconstructed from these angles and from the segment lengths. The segment lengths were measured on a frame taken in the middle of the stance phase for each limb. The distances between the torso marker and the hip and shoulder markers were measured on each frame to account for the movements of the hip and the shoulder relative to the torso (these distance *vs* time curves were smoothed using a spline function). The positions of the segments on the left side of the body were reconstructed on the assumption that the movements of the segments of one side of the body during a half stride were equal to those on the other side during the other half stride [23].

The position of the centre of mass and the moment of inertia about the centre of mass were calculated for each limb segment (Table S1), assuming each segment had the shape of a truncated cone and a uniform mass distribution (see Supplementary Materials). For the head and torso segments, the position of the centre of mass and the moment of inertia about the centre of mass (Table S1) were calculated assuming the segments had the shape of a sphere and a cylinder, respectively, with a uniform mass distribution. The circumference of each end and the length of each segment were measured on the elephants previous to the experiments. The proximal circumference of the thigh and the upper arm could not be measured on all the elephants, and were then set to 1.5 times the distal circumference of these segments; this ratio was approximated from measurements made on pictures and from data obtained on dissected segments by Hutchinson (unpublished data). Similarly, the radius of the sphere (head) was estimated on pictures, and was considered as 14% of the torso length. The segment masses (and the circumference of the torso) were estimated using volumes (and volumetric mass) reconstructed from scan and using the data of Ren and Hutchinson [24].

The horizontal and vertical position (*X, Y*) of the *COM* of the elephant was computed each frame by:

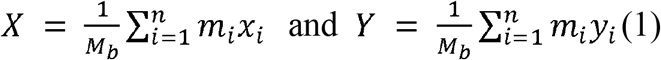

where *M*_b_ is the mass of the animal, and *m*_i_, *x*_i_, *y*_i_ are the mass, the horizontal, and the vertical position of the centre of mass of the *i*^th^ segment. The position of the centre of mass of each segment was obtained using the model of a frustum of a cone, or a cylinder for the trunk (see Supplementary Materials). Its position relative to the *COM* was computed as the difference between the absolute coordinates of the segments and those of the *COM*.

The angular velocity of each segment was calculated from the derivative of its angle *versus* time relationship. The relative linear velocity of each segment was calculated from the time-derivative of the coordinates of its centre of mass relative to the *COM*. The kinetic energy 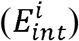 of the i^th^ segment was calculated as the sum of its translational and rotational energy:

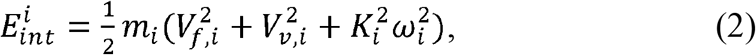

where *m*_i_ and *K*_i_ are the mass and radius of gyration of the i^th^ segment, *V*_f,i_ and *V*_v,i_ are the horizontal and vertical velocity of its centre of mass relative to the *COM* and ω_i_ is its angular velocity.

The energy-time curve of each limb was obtained by summing the energy-time curves of each of its segments. This procedure allowed energy transfer between the segments of the same limb.

The work done to move the hindlimbs 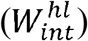, the forelimbs 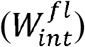, the head 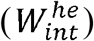 and the torso 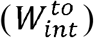 relative to the *COM* was obtained by summing the increments of the respective energy-time curves. The increments were considered to represent work actually done by the muscles only if the time between two successive maxima was greater than 10-80 ms, depending on the speed of progression. The total internal work (*W*_int_) was computed as:

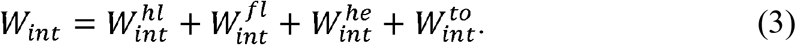

This equation allows energy transfers within the limbs but excludes transfers between the limbs, torso and/or head [8,25,26].

The stride/step selection was not made synchronously between *W*_int_ and *W*_ext_: a stride for internal work measurement was made between two successive right hindlimb touchdowns, and the selection of a stride/step for external work was made between peaks of velocity of the *COM* (see methods in Genin et al. [13] for more details). As a consequence, the total work (Eq. 4) is not computed point by point but by summing the average curves of *W*_int_ and *W*_ext_.

## Results

### Internal energy of the body segments

The *E*_int_ *versus* time curves of the limb-segments are shown in Fig. 2. The stick elephants on the bottom of the figure shows the position of the limbs every 12.5% of the stride. The mass of the hindfoot and forefoot each represent less than 1% of *M*_b_ whereas the shank and the forearm are more than twice as massive. Nevertheless, the amplitudes of their internal energy curves are similar because the two distal segments have a higher speed relative to the *COM* than the two intermediate segments. Note also that, despite their heavier mass (about 5% of *M*_b_), the amplitudes of the internal energy curves of the thigh and upper arm are smaller than the other four limb segments, in this case because they have a lower speed relative to the *COM*. Fig. 2 also shows that the energy curves of the segments of the same limb are for the most part in-phase, and as a consequence the amount energy that can be transferred from one segment to the other is rather small.

**Figure 2:**
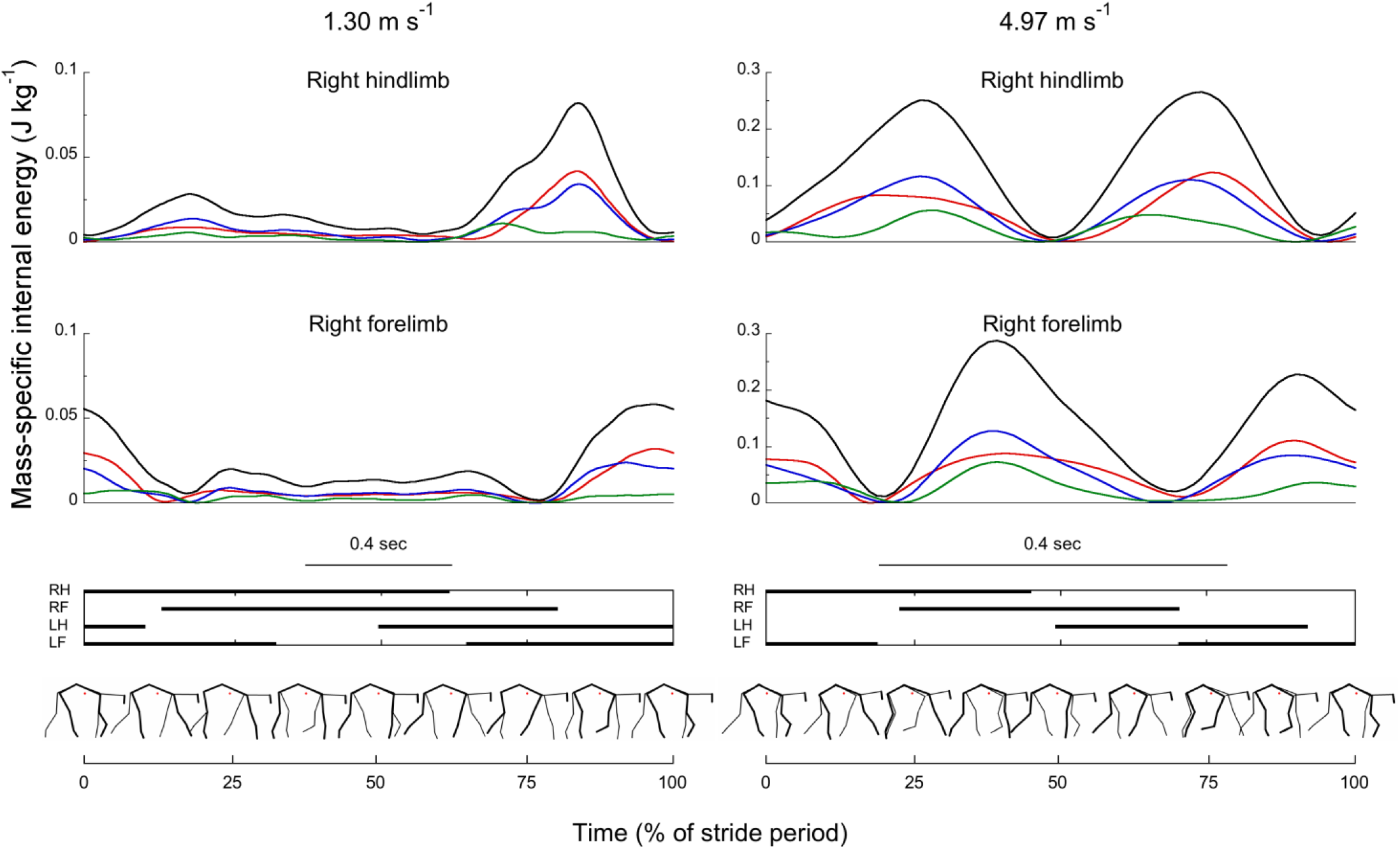
Typical traces of mass-specific kinetic energy curves of segments within the right hind and fore limbs due to their velocity relative to the *COM* during one stride, for an elephant of 1740 kg. Curves start and stop on successive touchdowns of the right hindlimb. Left curves are for a regular walking speed (1.3 m s^-1^) and right curves are for the fastest speed obtained (4.97 m s^-1^). These trials are similar to those used in Genin et al. (2009) to illustrate typical energy curves of the movement of the *COM* relative to the environment. The top pair of panels is for the segments of the right hindlimb, and the second pair of panels is for the segments of the right forelimb. The total kinetic energy of the limb (black curve) is the sum of the kinetic energy of the distal (red), intermediate (blue) and proximal (green) segments within the limb. The third panel indicates the footfall pattern of the elephant. The bottom panel shows stick elephants illustrating the position of the limbs every 12.5% of the stride cycle. At intermediate speed, the kinetic energy remains low when the limb touches the ground, and large oscillations occur during the swing of the limb. Hind and fore energy curves are not in phase due to the limb phase of ∼18% at that speed (see (Hutchinson et al., 2006)). At high speed, large oscillations of the kinetic energy occur during both the stance and the swing of the limb. Hind and forelimb energy curves are out of phase due to the limb phase of ∼23%. Note that for both speeds and both limbs, the small distal segment of the limb accounts for the largest part of the fluctuations of the energy curve of the total limb. A different scale is used between left and right side to draw attention to the influence of each segment on the limb energy curve.

Fig. 3 represents the time evolution of the internal energy curves of the hind and forelimbs, the torso and the head at intermediate and high speed of locomotion. Over one stride, the internal energy of the limbs presents two maxima, one when the limbs are moving forwards and a second when the limbs are moving backwards, relative to the *COM*. The torso and the head present four maxima over the stride, corresponding to the peak upwards and peak downwards speed of these segments relative to *COM* each step. The amplitude of the energy-time curves of the hind and forelimbs is much greater than that of the torso and of the head.

**Figure 3:**
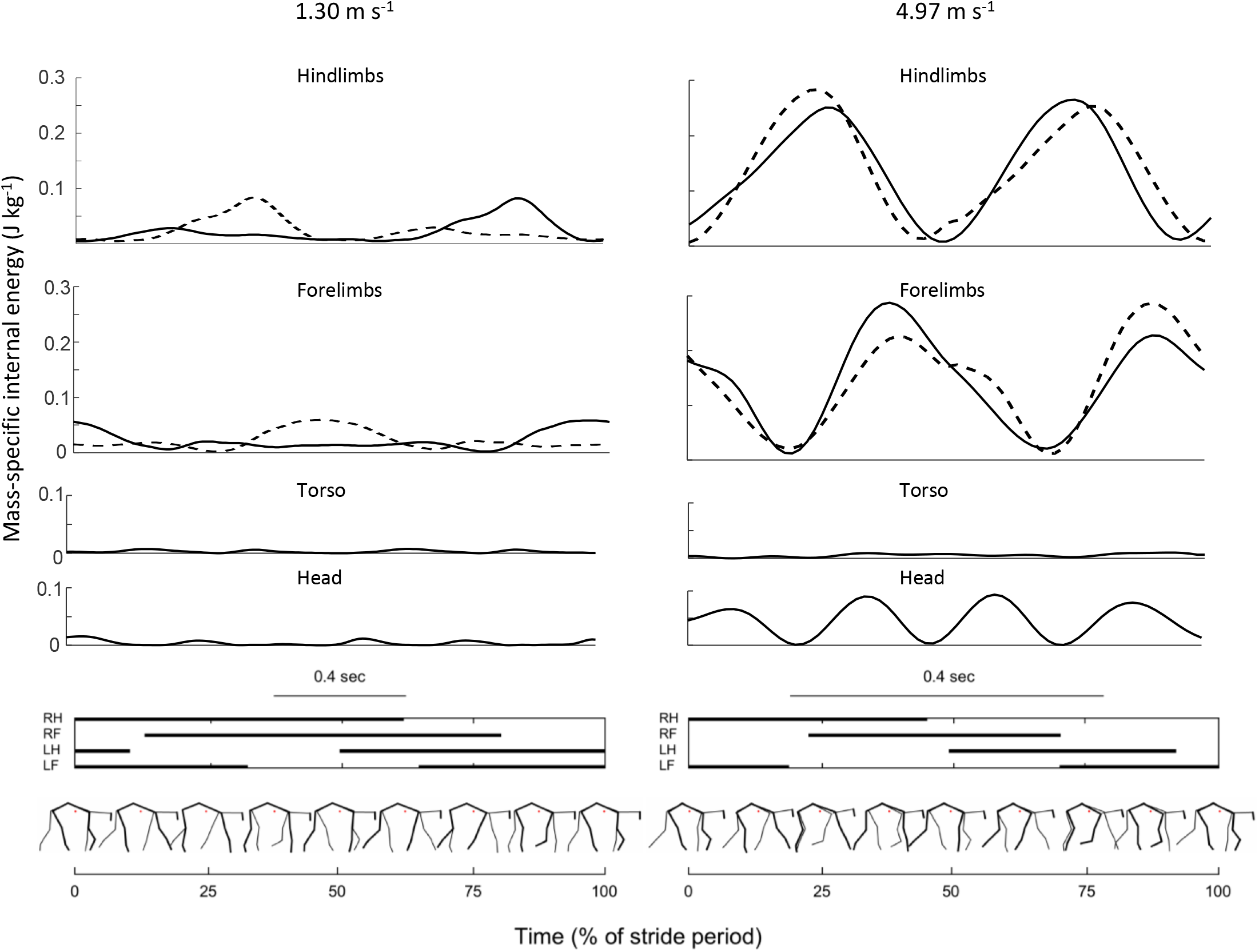
Effect of speed on typical traces of mass-specific kinetic energy curves of the limbs, the torso and the head due to their velocity relative to the *COM* during one stride. Thick lines indicate the position and kinetic energy of the right limbs. Energy curves for the left side look similar to the right side, as their values are equivalent to the right side, shifted by 180° of a stride period (see Methods for more details). Energy oscillations of the limbs increase greatly between intermediate and high speed. Internal energy of the torso and the head segments (55 and 16% of *M*_b_ respectively) remain almost nil at intermediate speed, whereas the energy of the head segment shows larger oscillations at maximal speed. This illustrates the importance of the head segment in our model. Other details similar to Fig. 2.

At low speeds, the period when a limb is moving backwards relative to the *COM* (∼contact period) is longer than the period when it is moving forwards (∼swing period). Consequently, its velocity and its internal energy are smaller when the limb moves backwards than when it swings forwards (Fig. 3 left). When speed increases, the relative contact period of each limb is reduced and as a consequence, the energy variations of the limbs during the contact phase and during the swing phase are about equal (Fig. 3 right).

### Internal work done to move the limbs relative to the COM

The mass-specific internal work per unit distance (J kg^-1^ m^-1^) done to move the two Hindlimbs 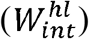, the two forelimbs 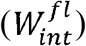, the head 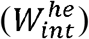 and the torso 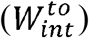 relative to the *COM* is presented in Fig. 4A, together with the total internal work (*W*_int_), which is the sum of 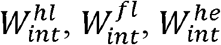 and 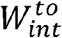. The *W*_int_ presents a curvilinear increase with the speed of locomotion: from ∼0.1 J kg^-1^ m^-1^ at low speeds to ∼0.7 J kg^-1^ m^-1^ at the highest speeds.

**Figure 4.**
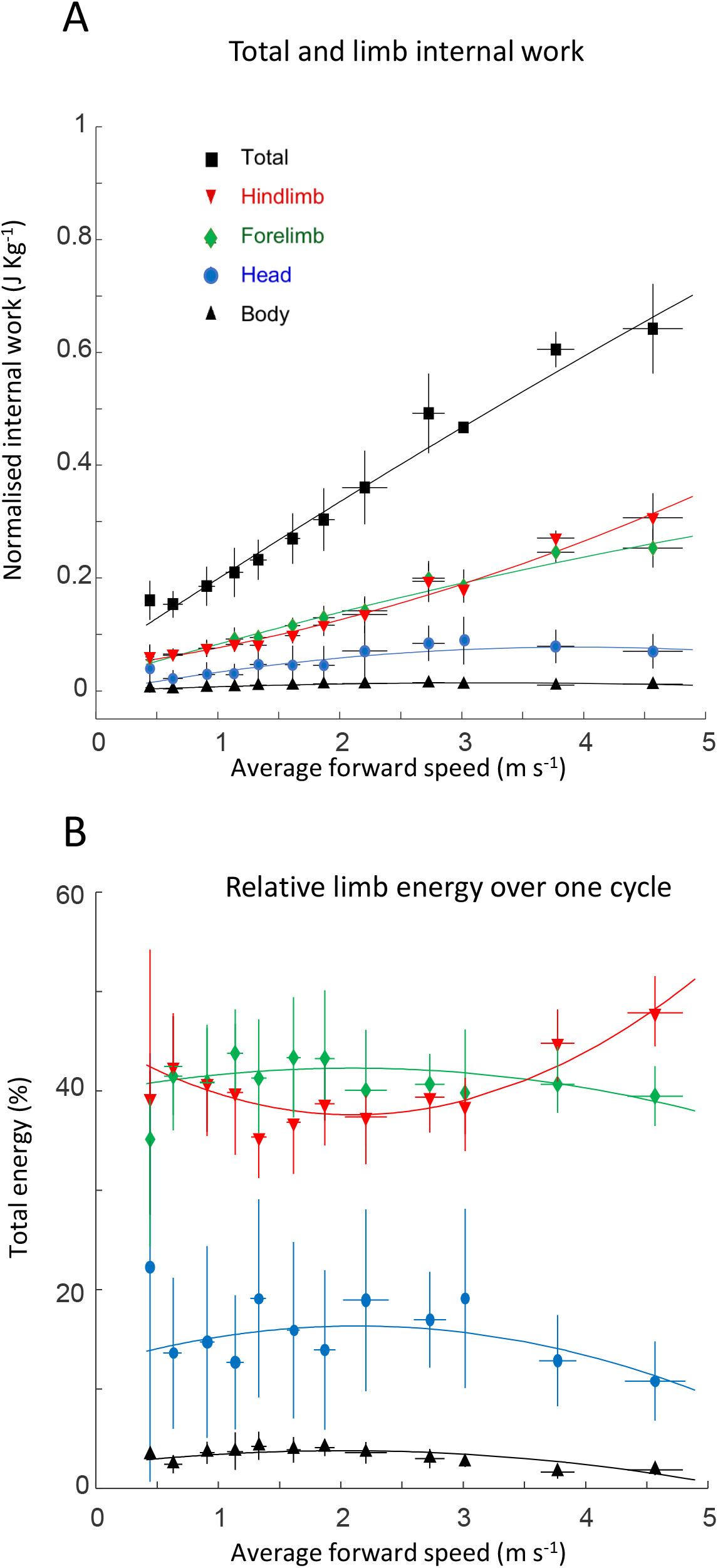
(A) Effect of speed on the mass-specific positive work per unit-distance associated with the kinetic energy changes due to the movement of all the body segments relative to the *COM* (*W*_int_, black squares), computed as the sum of the internal positive work of the two hind (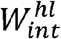, red triangle) and the forelimbs (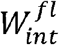, green diamond), the torso (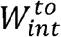, black triangles) and the head (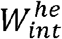, blue circles). Lines are second order polynomial curve fits. The most representative curve fit of *W*_int_ (in J kg^-1^ m^-1^) vs speed (*V*_f_ in m s^-1^) is a quadratic function of 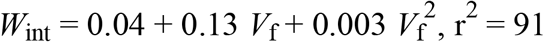. (B) The internal work due to hind and front limbs, the head and the torso as a fraction of the total internal work. 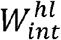 and 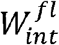 each account for an average 40-45% of *W*_int._. The internal work due to the movements of the head, 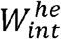, accounts for ∼14% of *W*_int_ and is mainly due to the vertical movements of the head relative to the *COM* (the horizontal distance between the *COM* and the head doesn’t change much, and the rotational movement of the head was not measured here).

As speed increases, both the step length and the step frequency increase (see Figs. 1 & 2 in Genin et al., [13]). As a consequence, the speed of the limb segments relative to the *COM* and thus the work 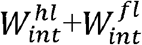 becomes more important (Fig. 4A, green and red symbols). Although the mass of the four limbs is less than 30% of *M*_b_, the work done to move these segments represents about 80% of *W*_int_ at 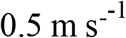 and about 95% at 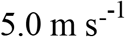 (Fig. 4B). As speed increases the fraction of *W*_int_ due to movements the limbs 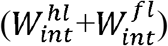 is slightly increased. This is due to the relative decrease in the work done to bob the head 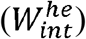, which decreases from ∼15% of *W*_int_ at low speeds to about 10% of *W*_int_ at the highest speed. At all speeds the work done to move the torso represents less than 5% of the *W*_int_, despite the fact that the mass of the torso represents ∼55% of *M*_b_.

## Discussion

The internal work (*W*_int_) as measured in this study includes only the work done to accelerate the body segments relative to the *COM*. There are many other types of internal work; in fact, internal work is sufficiently complex that it is easier to define what it is not. Internal work is not any of the work done against the environment, such as work done against the resistance of air or done to move the substrate under the feet [27–29]; neither is internal work any of the work done to lift or accelerate the *COM* on slopes [22,30–32]-these are all external work. However, internal work is all of the rest of the work done during locomotion. For example, not counted in this study, internal work is the work done to pump air, blood, and calcium; it is the work done by one muscle against another during antagonistic contractions; it is the work done by one leg against another during a step phase involving multiple foot contacts [33]. In short, it is all the work done that does not result in a displacement of the *COM* or the environment.

Some of these other types of internal work are not insignificant. For example, the work done by one leg against another during ground contact, *W*_int, dc_, not included here, accounts for up to 10% of the total work in humans at intermediate walking speeds, although it decreases rapidly as walking speed is either increased or decreased [25,33]. Also in elephants, limbs generated or absorbed a substantial amount of simultaneous positive and negative power during double or triple support [14]. However, in their ‘individual limb methods’ analysis, Ren et al. [14] summed the power curve of all limbs, allowing transfer between them (Fig. 5A). Such transfer of energy may allow a potential compression and rebound of each limb during the stance (and thus a potential elastic energy storage and release), without increasing the vertical displacement of the COM (and thus *W*_ext_) [14,34]. This is made possible by rotation of the trunk (Fig. 5B), which increases until about 2 m s^-1^ and decreases at higher speeds. Contrary to the increase in *W*_int_ with increasing speed, showing no discontinuity, the changes in trunk rotation may indicate that elephants use different gaits throughout their entire speed range.

**Figure 5.**
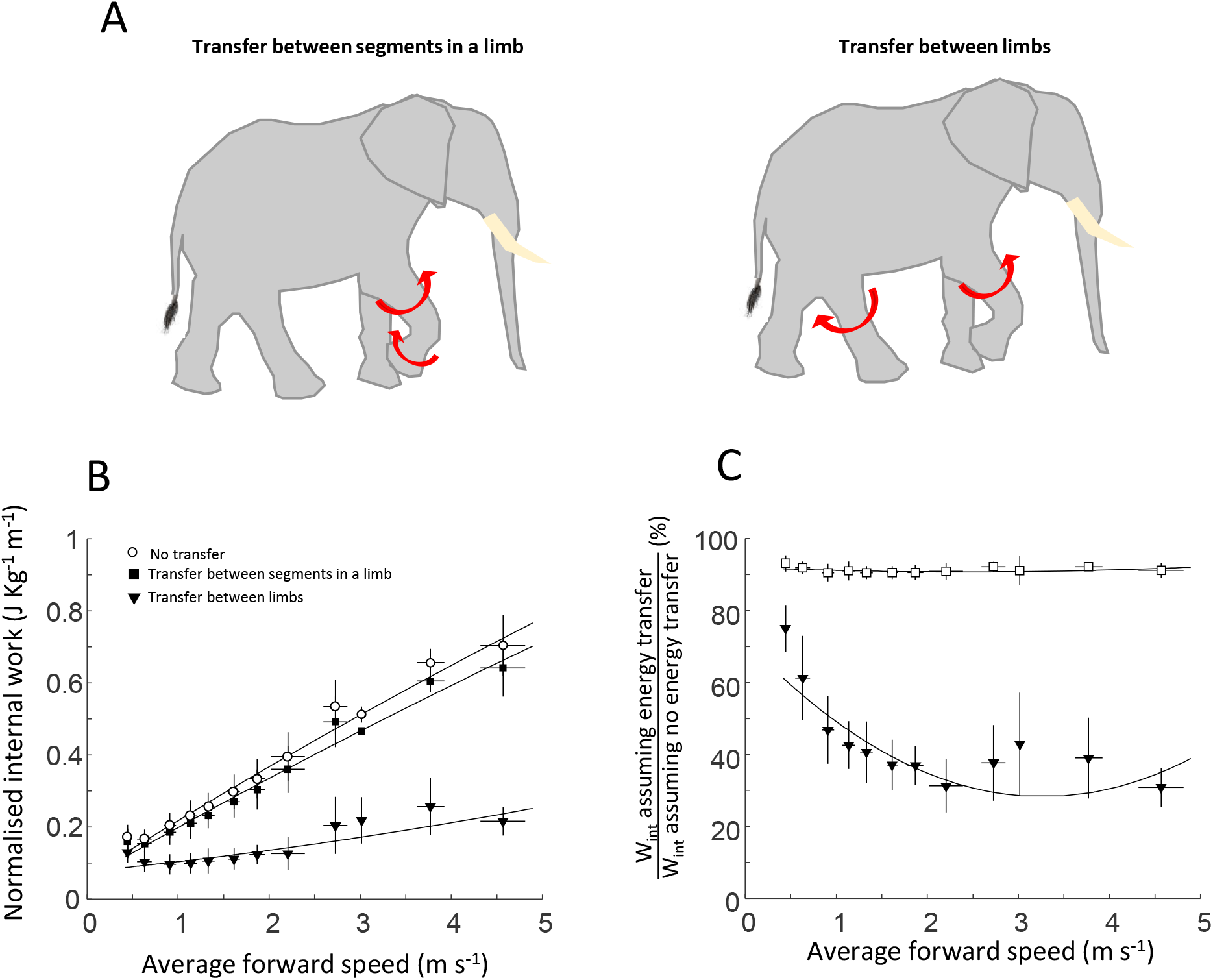
(A) Effect of energy transfer on the mass-specific internal work per unit distance. The initial values for internal work (*W*_int_, from Fig. 4, black squares) include energy transfer between segments within a limb, but not between segments of two different limbs. In order to assess the effect of energy transfers, the internal work excluding any energy transfer (*W*_int, max_, circles) and including energy transfer across the trunk (*W*_int, min_, triangles) are shown (Willems et al., 1995). (B) *W*_int_ and *W*_int, min_ expressed as a fraction of the upper limit for internal work, *W*_int, max_. Including energy transfer between the segments within each limb in our calculation on elephants reduces *W*_int_ independently of speed by no more than 10%. Assuming energy transfer between all the segments within each limb, the head and torso (open triangles) results in a reduction of *W*_int_ by up to 70% at intermediate and fast speeds.

Recently, it has been shown that the mechanical energy fluctuations due to soft tissue energy dissipation (heel pad and foot arch compressions, visceral sway, cartilage and intervertebral disc compressions etc.) can also affect the mechanical work production in locomotion [35–37]. During walking, the body’s soft tissues deform and dissipate a fraction of the mechanical energy (negative mechanical work) that must be actively replaced by the muscles to maintain a constant speed (positive mechanical work) [35]. The energy dissipated by the soft tissues can be estimated by comparing different methods used to calculate the mechanical energy output of the body [38]. Therefore, the difference between both methods represents, among others, the amount of energy being dissipated by the soft tissues [35,39,40]. The discrepancy between our mechanical work measurements and the data published by Ren et al. [14] may partly be due to soft tissue vibration, increasing with speed (Fig 5C), as in human running [36]. In any case, the work done to move the body segments relative to the *COM* is by far the greatest single type of internal work during normal-legged terrestrial locomotion.

### Maximal and minimal limits of internal work

Cavagna and Kaneko measured the mechanical work required to move the limbs relative to the *COM* in human walking and running and discussed the different possibilities of energy transfer between different segments of the body [41]. Willems et al. [8] have done the most detailed analysis to date on the possibility of energy transfers between segments as well as between the segments and the *COM* of the whole body. Indeed, the possibility exists that, when the energy of one segment (or the *COM*) decreases simultaneously with an increase of a different segment (or the *COM*), the measured increase in energy is due to a passive transfer of energy rather than due to the performance of muscular work (see simplified examples in Fig. 6A). In this case *W*_int_, measured as the sum of all the increases in kinetic energy for each segment taken individually, would be greater than the work done by the muscles.

**Figure 6.**
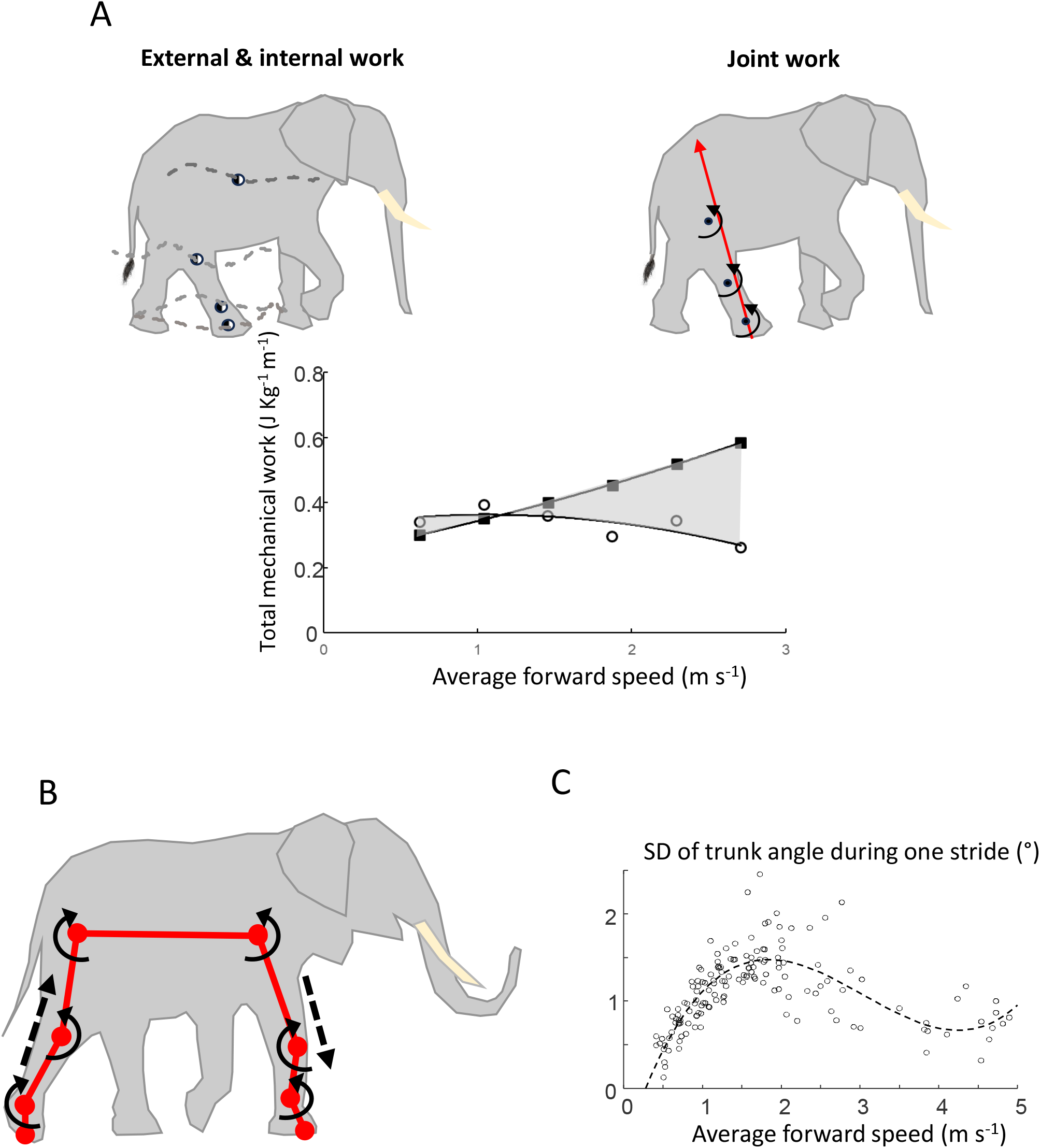
The mass-specific internal (*W*_int_, panel D), external (*W*_ext_, panel C) and total (*W*_tot_, panel B) mechanical work per unit distance are represented as a function of the speed of locomotion in elephants (thick lines) and walking (W) and trotting (T) horses (thin lines). The external work in elephants shows a U-curve and a slight minimum at intermediate ‘optimal’, speed of locomotion. *W*_int_ is greater than *W*_ext_ at all except the very lowest speeds. *W*_int_ and *W*_ext_ in elephants show a continuous evolution throughout the entire range of speed, whereas the gait transition from a walk to a trot in horse shows a clear difference in the work value and the rate of the work as a function of speed, for internal, external and total work. The value for total work as a function of speed, *W*_tot_, is computed by summing the average curves of *W*_int_ and *W*_ext_ (Eq. 5). Metabolic cost of locomotion (dotted lines, B) show a U-shape curve for both elephants and walking or trotting horses. The locomotory efficiency (panel A), computed as the ratio between total mechanical work and the metabolic cost, shows an inverted U-shaped curve with a maximum of 45% at 1.24 m s^-1^ in elephant.

An intersegment transfer of kinetic energy may take place (i) across the trunk (*e*.*g*. from the right hindlimb to the left forelimb; Fig. 5A) or (ii) between segments of a limb (*e*.*g*. from the thigh to the shank; Fig. 6A-left). The two previous studies [8,41] concluded that the most accurate computation was to include only energy transfers between segments within a limb, and to exclude transfers between segments of two different limbs; this is the method used to calculate *W*_int_ in this study. In other words, the value of the internal work in elephants as a function of the speed of progression shown in Fig. 4A was determined by adding at each instant the kinetic energy of the three segments of each limb, and then summing the increases of the resultant curve (shown in Fig. 3) to determine the positive muscular work done.

To assess the maximum influence of other possible, if unlikely, energy transfers on the value of the internal work, complementary computations of the internal work were made: (i) excluding any energy transfer (*W*_int, max_), resulting in a maximum limit for the measure of internal work; and (ii) including all energy transfers, including transfers across the trunk (*W*_int, min_), representing the minimum value for *W*_int_ (however not including energy transfers between *W*_int_ and *W*_ext_). The *W*_int, max_ was computed by summing the positive increments of all 14 segment energy curves, while *W*_int, min_ was obtained by summing all 14 segment energy curves and then measuring the positive increments of the single resulting energy curve (Fig. 5).

Maximal and minimal limits of the muscular work required to move the limbs, the head and the trunk relative to the *COM* are represented as a function of speed in Fig. 5A for elephants, while the influence of energy transfers on the amplitude of *W*_int_ and *W*_int, min_ (respectively filled squares and open triangles) relative to *W*_int, max_ is shown in Fig. 5B. The upper line in Fig. 5B (squares) shows that allowing energy transfers between the segments within the limb reduces *W*_int_ independently of speed by no more than 10%, a result similar to that found in human walking or running [8]. On the other hand, allowing any energy transfers between all 14 of the body segments (open triangles) would result in a *W*_int_ reduction of up to 70% at intermediate and fast speeds, which is twice the maximum reduction found in humans [8]. Such a difference in maximum *W*_int_ reduction (*W*_int, min_, in Fig. 5A) between elephants and humans can be explained in part by the shift of ∼20% in the stride phase between ipsilateral limb touchdowns (see Fig. 2 of Genin et al., [13]), which is the possible occasion of large amounts of energy transfer (see right panels of Fig. 2). In contrast, the human ipsilateral upper and lower limbs oscillate almost simultaneously in opposite directions, but upper and lower limb energy curves increase and decrease together (note that kinetic energy is independent of the direction of the speed). This results in smaller possible energy transfers between limbs across the trunk and limits the decrease of *W*_int, min_ to ∼30% relative to *W*_int, max_ in humans (see Fig. 1, 7 and 8 in Willems et al., [8]).

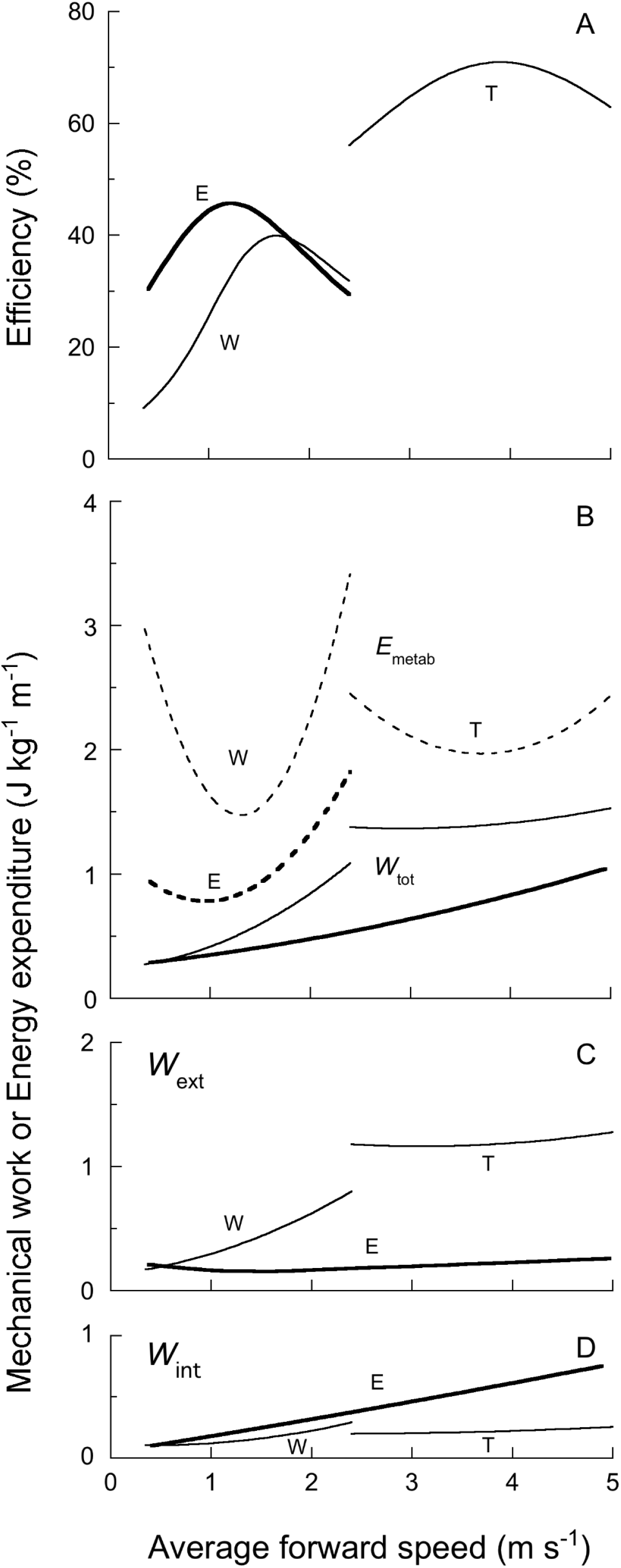

While the *W*_int, min_ represents the arithmetic minimum limit to the internal work done during elephant locomotion, we do not consider this limit to be realistic for the following reasons. First, there is no anatomical structure for transferring kinetic energy, for example from the forefoot to the hindfoot. Second, although kinetic energy is a scalar and therefore increases with increases in both positive and negative segment velocity changes relative to the *COM*, clearly kinetic energy transfers cannot occur between two segments which both undergo velocity changes in the same direction, as happens often in segments of ipsilateral limbs during elephant locomotion. Thus, we consider the most correct solution is to allow energy transfers (where possible) within limbs but not between limbs (as has been done previously, [8]); all the discussion that follows is predicated upon this assumption.

### Internal, external work and total work

The total muscle and tendon work required to maintain the energy fluctuations of the body during steady-state locomotion can be computed as:

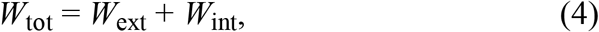

where *W*_ext_ is the mechanical work necessary to maintain the motion of the *COM* of the whole body during locomotion (*e*.*g*. in humans [22,30,42]; in non-human animals [11,13,43]), and *W*_int_ is the mechanical work necessary to maintain the movements of the body segments relative to the *COM* [8,9,11,26]. It has been shown in humans that energy transfers between *W*_ext_ and *W*_int_ are quite small [8], and we have assumed in Eq. 4 that any similar transfers in elephants are negligible.

Fig. 7 shows the mass -specific internal (panel D), external (panel C) and total (panel B) work per unit distance as a function of the speed of locomotion in elephants (thick lines) as compared to horses walking and trotting (thin lines, Minetti et al., [44]). The *W*_ext_ is almost independent of the speed, whereas *W*_int_ increases curvilinearly with speed. The curve fit of *W*_tot_ (in J kg^-1^ m-^-1^) versus speed (*V*_f_ in m s-^1^) is a quadratic function:

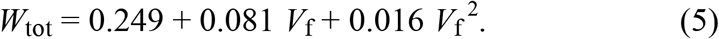

The ratio of *W*_int_ to *W*_ext_ shows their relative importance as a function of speed. In most animals, the ratio is considerably less than 1 at low speeds. However, since the *W*_ext_ remains roughly constant while *W*_int_ increases exponentially with speed, the ratio often approaches 2 at high speeds. Notably, this ratio attains values up to 3:1 in fast-moving elephants, a much higher ratio than seen in cursorial animals. Even heavy-legged humans, who attain much higher speeds than elephants, only reach a ratio of ∼2:1 [8]. The high *W*_int_:*W*_ext_ ratio in elephants is a result of heavy legs, a heavy head, an unusually high step frequency, and a relatively low *W*_ext_ [13]. In humans, at any given speed a high step frequency increases *W*_int_ and decreases *W*_ext_ [9,21].

The walk-trot gait transition in a horse shows a clear difference in the work value and the slope of the work as a function of speed, for internal, external and total work. These changes are typical indicators (amongst others) of a change in gait and can be observed in most species [10–12]. In contrast, internal, external and total work in elephants show a continuous evolution throughout their entire range of speed, suggesting that the gait transition is smoother than observed in other animals.

### Efficiency of positive work during elephant locomotion

Efficiency is one of the fundamental properties of any ‘energy transducer’, something that converts energy from one form to another. During locomotion, all animals convert metabolic energy into mechanical work. In this case, the efficiency is the ratio between the mechanical work performed and the metabolic energy consumed to do that work. Efficiency has no units since both the numerator and the denominator are units of energy (or energy/time). The main conceptual difficulty of determining the efficiency of animal locomotion comes from the definition of the work of locomotion. In this study, we have taken the approach of many before us, and have indirectly measured the net positive muscular work while acknowledging that part of this positive work may have been done by tendons and other elastic structures. Any increase in the total kinetic and gravitational potential energy of the body is assumed to be due to positive muscular work (in reality, part of it comes from the energy stored in elastic structures); any decrease in the total kinetic and gravitational potential energy of the body is assumed to be lost as heat (in reality, part of it is stored in elastic structures).

To a lesser extent, metabolic energy consumption also presents conceptual problems. Although it is generally agreed that steady-state oxygen consumption is a good measure of energy input to a subject, much of animal locomotion occurs at speeds that are not steady state. Furthermore, there is the question of whether the basal, resting or standing metabolism (or nothing) should be subtracted from the steady-state value when calculating the efficiency.

The metabolic energy expenditure during locomotion minus standing has been measured in a previous study [4] in three young African elephants (average mass 1542 kg) at speeds up to 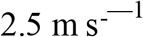, and is shown in Fig. 7B (thick dotted line), along with data from horses for comparison to the next-largest animal with comparable data (thin dotted lines, Minetti et al., [44]). Although our work measurements were conducted on Asian elephants, various studies have shown no differences between African and Asian elephants in kinematic parameters [24,45] or energy changes of the *COM* as measured by motion sensors [24].

The efficiency is shown in Fig. 6A for both the elephant (thick line) and horse walk and trot (thin lines). Efficiency is computed as the ratio between total mechanical work (plain lines panel B) and the net metabolic cost (dotted lines panel B) in elephants (thick lines) and walking (W) and trotting (T) horses (thin lines). Efficiency in elephants was calculated over the speed range that metabolic data is available (up to 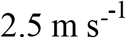), and shows an inverted U-shape curve with a maximum of 45% at 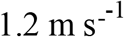. The optimal efficiency occurs close to the speed where the pendular mechanism of walking is optimal (reflected by the percent recovery in Genin et al., [13] Fig. 4A), and consequently where the external mechanical work is minimal [43], although this would tend to minimise the efficiency if not for the increase in internal work with speed coupled with a strong minimum in the energy expenditure. A peak walking efficiency of 45% in walking elephants compares favourably with that in the horse (Fig. 6A) and in humans [8,43], lending credence to the notion that elephants are very competent walkers at low speeds [13].

Assuming the maximum efficiency of muscle shortening without a prestretch is about 25% [46], an efficiency of locomotion >25% indicates that part of the work done during locomotion must come either from enhanced non-elastic contractility of muscle [47] or energy stored in elastic elements during a previous negative work phase of the step [21].

### Mechanical work and body size

In the early ‘80s, Taylor’s group at Harvard’s Concord Field Station published a series of papers on the mechanical work and metabolic cost of locomotion in animals of different sizes. They concluded that the mass-specific metabolic power (Ė_metab_, in W kg^-1^) increased linearly with the speed of progression, but at a given speed, decreased with body size over a 4-orders of magnitude size range from 0.01 to 260 kg [2]. In comparison, the mass-specific total mechanical power (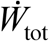, calculated in the same manner as in this study, in 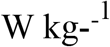) increased curvilinearly with speed of progression but appeared to be independent of body size over a 3-orders of magnitude size range from 0.043 to 70 kg [10–12].

This led to some speculation in the literature involving extrapolation of the mechanical power data to animals the size of adult elephants, with the conclusion that the two lines cross at a body size of about 2 tons [48]. In other words, extrapolating from animals less than 70 Kg to animals larger than 2 tons leads to the unlikely muscular efficiency calculation of more than 100%. Although this value is not impossible given the way the mechanical power is measured (as noted above, all work done by elastic structures is attributed to the muscles), it would require that very large animals be extremely ‘bouncy’ – not a characteristic normally attributed to adult elephants [17,49].

Since the original Harvard studies, metabolic data on very large animals have become available, and the up to 3500 kg elephants [4,5] show a reasonably good fit to an extrapolation of the allometric equation of Taylor et al. [2]. The current study provides the first data on very large animals that can be compared to the total mechanical work and efficiency measurements of the Harvard studies [12]. It appears that the allometric equation of [12] relating total mechanical power output to body size, based upon just 5 species and a limited size range, cannot be used to accurately predict power outputs for animals that are ∼50 times larger than the animals measured. The mechanical power data of Minetti et al. [44] on horses supports this conclusion as well. Unfortunately, the too-high mechanical power predictions have led to erroneous conclusions of very high efficiency in large animals [3,48]. In fact, the measured efficiency in walking elephants is quite similar to that in much smaller horses (Fig. 6A) and humans.

## Notes

### Competing Interest Statement

The authors have declared no competing interest.

## REFERENCES

1. Clemente CJ, Dick TJM. 2023 How scaling approaches can reveal fundamental principles in physiology and biomechanics. Journal of Experimental Biology 226. (doi:10.1242/jeb.245310)

2. Taylor CR, Heglund NC, Maloiy GM. 1982 Energetics and mechanics of terrestrial locomotion. I. Metabolic energy consumption as a function of speed and body size in birds and mammals. J Exp Biol 97, 1–21. (doi:10.1242/jeb.97.1.1)

3. Full R, Wieser W, Gnaiger A. 1989 Mechanics and energetics of terrestrial locomotion: bipeds to polypeds. Energy transformations in cells and organisms., 175–182.

4. Langman VA, Roberts TJ, Black J, Maloiy GM, Heglund NC, Weber J-M, Kram R, Taylor CR. 1995 Moving cheaply: Energetics of walking in the African elephant. Journal of Experimental Biology 198, 629–632.

5. Langman VA, Rowe MF, Roberts TJ, Langman NV, Taylor CR. 2012 Minimum cost of transport in Asian elephants: Do we really need a bigger elephant? Journal of Experimental Biology 215, 1509–1514. (doi:10.1242/jeb.063032)

6. Peyré-Tartaruga LA et al. 2021 Mechanical work as a (key) determinant of energy cost in human locomotion: recent findings and future directions. Exp Physiol 106, 1897–1908. (doi:10.1113/EP089313)

7. Heglund NC. 2004 Physiology. Running a-fowl of the law. Science 303, 47–48. (doi:10.1126/science.1093588)

8. Willems PA, Cavagna GA, Heglund NC. 1995 External, internal and total work in human locomotion. J. Exp. Biol. 198, 379–393.

9. Mesquita RM, Willems PA, Catavitello G, Dewolf AH. 2023 Kinematics and mechanical changes with step frequency at different running speeds. Eur J Appl Physiol (doi:10.1007/s00421-023-05303-3)

10. Fedak MA, Heglund NC, Taylor CR. 1982 Energetics and mechanics of terrestrial locomotion. II. Kinetic energy changes of the limbs and body as a function of speed and body size in birds and mammals. J Exp Biol 97, 23–40. (doi:10.1242/jeb.97.1.23)

11. Heglund NC, Cavagna GA, Taylor CR. 1982 Energetics and mechanics of terrestrial locomotion. III. Energy changes of the centre of mass as a function of speed and body size in birds and mammals. J Exp Biol 97, 41–56. (doi:10.1242/jeb.97.1.41)

12. Heglund NC, Fedak MA, Taylor CR, Cavagna GA. 1982 Energetics and mechanics of terrestrial locomotion. IV. Total mechanical energy changes as a function of speed and body size in birds and mammals. Journal of Experimental Biology 97, 57–66.

13. Genin JJ, Willems PA, Cavagna GA, Lair R, Heglund NC. 2010 Biomechanics of locomotion in Asian elephants. J. Exp. Biol. 213, 694–706. (doi:10.1242/jeb.035436)

14. Ren L, Miller CE, Lair R, Hutchinson JR. 2010 Integration of biomechanical compliance, leverage, and power in elephant limbs. Proc Natl Acad Sci U S A 107, 7078–7082. (doi:10.1073/pnas.0911396107)

15. Hill AV. 1950 The Dimensions of Animals and Their Muscular Dynamics. Science Progress (1933-) 38, 209–230.

16. Taylor CR, Shkolnik A, Dmi’el R, Baharav D, Borut A. 1974 Running in cheetahs, gazelles, and goats: energy cost and limb configuration. Am J Physiol 227, 848–850. (doi:10.1152/ajplegacy.1974.227.4.848)

17. Kram R, Taylor CR. 1990 Energetics of running: a new perspective. Nature 346, 265–267. (doi:10.1038/346265a0)

18. Marsh RL, Ellerby DJ, Carr JA, Henry HT, Buchanan CI. 2004 Partitioning the energetics of walking and running: swinging the limbs is expensive. Science 303, 80–83. (doi:10.1126/science.1090704)

19. Hildebrand M, Hurley JP. 1985 Energy of the oscillating legs of a fast-moving cheetah, pronghorn, jackrabbit, and elephant. Journal of Morphology 184, 23–31. (doi:10.1002/jmor.1051840103)

20. Cavagna GA, Willems PA, Franzetti P, Detrembleur C. 1991 The two power limits conditioning step frequency in human running. J Physiol 437, 95–108. (doi:10.1113/jphysiol.1991.sp018586)

21. Cavagna GA, Franzetti P, Heglund NC, Willems P. 1988 The determinants of the step frequency in running, trotting and hopping in man and other vertebrates. J. Physiol. (Lond.) 399, 81–92.

22. Dewolf AH, Peñailillo LE, Willems PA. 2016 The rebound of the body during uphill and downhill running at different speeds. Journal of Experimental Biology 219, 2276–2288. (doi:10.1242/jeb.142976)

23. Dewolf AH, Ivanenko Y, Zelik KE, Lacquaniti F, Willems PA. 2018 Kinematic patterns while walking on a slope at different speeds. J Appl Physiol (1985) 125, 642–653. (doi:10.1152/japplphysiol.01020.2017)

24. Ren L, Hutchinson JR. 2008 The three-dimensional locomotor dynamics of African (Loxodonta africana) and Asian (Elephas maximus) elephants reveal a smooth gait transition at moderate speed. Journal of the Royal Society Interface 5, 195–211. (doi:10.1098/rsif.2007.1095)

25. Schepens B, Bastien GJ, Heglund NC, Willems PA. 2004 Mechanical work and muscular efficiency in walking children. Journal of Experimental Biology 207, 587–596. (doi:10.1242/jeb.00793)

26. Dewolf AH, Ivanenko YP, Zelik KE, Lacquaniti F, Willems PA. 2019 Differential activation of lumbar and sacral motor pools during walking at different speeds and slopes. J. Neurophysiol. 122, 872–887. (doi:10.1152/jn.00167.2019)

27. Lejeune TM, Willems PA, Heglund NC. 1998 Mechanics and energetics of human locomotion on sand. J. Exp. Biol. 201, 2071–2080.

28. Dewolf AH, Lejeune TM, Willems PA. 2019 The on-off ground asymmetry during running on sand. Computer Methods in Biomechanics and Biomedical Engineering 22(supp 1), 325–327.

29. Pugh LG. 1971 The influence of wind resistance in running and walking and the mechanical efficiency of work against horizontal or vertical forces. J. Physiol. (Lond.) 213, 255–276. (doi:10.1113/jphysiol.1971.sp009381)

30. Dewolf AH, Ivanenko YP, Lacquaniti F, Willems PA. 2017 Pendular energy transduction within the step during human walking on slopes at different speeds. PLoS ONE 12, e0186963. (doi:10.1371/journal.pone.0186963)

31. Gomeñuka NA, Bona RL, da Rosa RG, Peyré-Tartaruga LA. 2016 The pendular mechanism does not determine the optimal speed of loaded walking on gradients. Hum Mov Sci 47, 175–185. (doi:10.1016/j.humov.2016.03.008)

32. Minetti AE, Ardigò LP, Saibene F. 1993 Mechanical determinants of gradient walking energetics in man. J Physiol 472, 725–735. (doi:10.1113/jphysiol.1993.sp019969)

33. Bastien GJ, Heglund NC, Schepens B. 2003 The double contact phase in walking children. Journal of Experimental Biology 206, 2967–2978. (doi:10.1242/jeb.00494)

34. Lambert A-S, Genin J, Heglund NC. 2010 A simple spring-mass model of running does not apply to elephants. Computer Methods in Biomechanics and Biomedical Engineering 13, 81–82. (doi:10.1080/10255842.2010.493732)

35. Riddick RC, Kuo AD. 2022 Mechanical work accounts for most of the energetic cost in human running. Sci Rep 12, 645. (doi:10.1038/s41598-021-04215-6)

36. Riddick RC, Kuo AD. 2016 Soft tissues store and return mechanical energy in human running. J Biomech 49, 436–441. (doi:10.1016/j.jbiomech.2016.01.001)

37. Zelik KE, Kuo AD. 2010 Human walking isn’t all hard work: evidence of soft tissue contributions to energy dissipation and return. J Exp Biol 213, 4257–4264. (doi:10.1242/jeb.044297)

38. Winter DA. 1979 Biomechanics of human movement. Wiley.

39. Fu X-Y, Zelik KE, Board WJ, Browning RC, Kuo AD. 2015 Soft Tissue Deformations Contribute to the Mechanics of Walking in Obese Adults. Med Sci Sports Exerc 47, 1435–1443. (doi:10.1249/MSS.0000000000000554)

40. Zelik KE, Kuo AD. 2012 Mechanical work as an indirect measure of subjective costs influencing human movement. PLoS One 7, e31143. (doi:10.1371/journal.pone.0031143)

41. Cavagna GA, Kaneko M. 1977 Mechanical work and efficiency in level walking and running. J Physiol 268, 467--81. (doi:10.1113/jphysiol.1977.sp011866)

42. Cavagna GA, Thys H, Zamboni A. 1976 The sources of external work in level walking and running. J. Physiol. (Lond.) 262, 639–657. (doi:10.1113/jphysiol.1976.sp011613)

43. Cavagna GA, Heglund NC, Taylor CR. 1977 Mechanical work in terrestrial locomotion: two basic mechanisms for minimizing energy expenditure. Am. J. Physiol. 233, R243–261.

44. Minetti AE, Ardigò LP, Reinach E, Saibene F. 1999 The relationship between mechanical work and energy expenditure of locomotion in horses. Journal of Experimental Biology 202, 2329–2338.

45. Hutchinson JR, Schwerda D, Famini DJ, Dale RHI, Fischer MS, Kram R. 2006 The locomotor kinematics of Asian and African elephants: changes with speed and size. J Exp Biol 209, 3812–3827. (doi:10.1242/jeb.02443)

46. Hill AV. 1964 The efficiency of mechanical power development during muscular. Proceedings of the Royal Society of London. Series B, Containing papers 159, 319–324. (doi:10.1098/rspb.1964.0005)

47. Mantovani M, Heglund NC, Cavagna GA. 2001 Energy transfer during stress relaxation of contracting frog muscle fibres. Journal of Physiology 537, 923–939. (doi:10.1113/jphysiol.2001.012657)

48. Alexander RM. 2005 Models and the scaling of energy costs for locomotion. Journal of Experimental Biology 208, 1645–1652. (doi:10.1242/jeb.01484)

49. Taylor CR. 1985 Force development during sustained locomotion: A determinant of gait, speed and metabolic power. Journal of Experimental Biology VOL. 115, 253–262.

