## Supplementary Materials for "Locomotion Efficiency of Elephants: Mechanical work and energetics"

**Supporting informations**

**Model of a frustrum of a cone and cylinder**

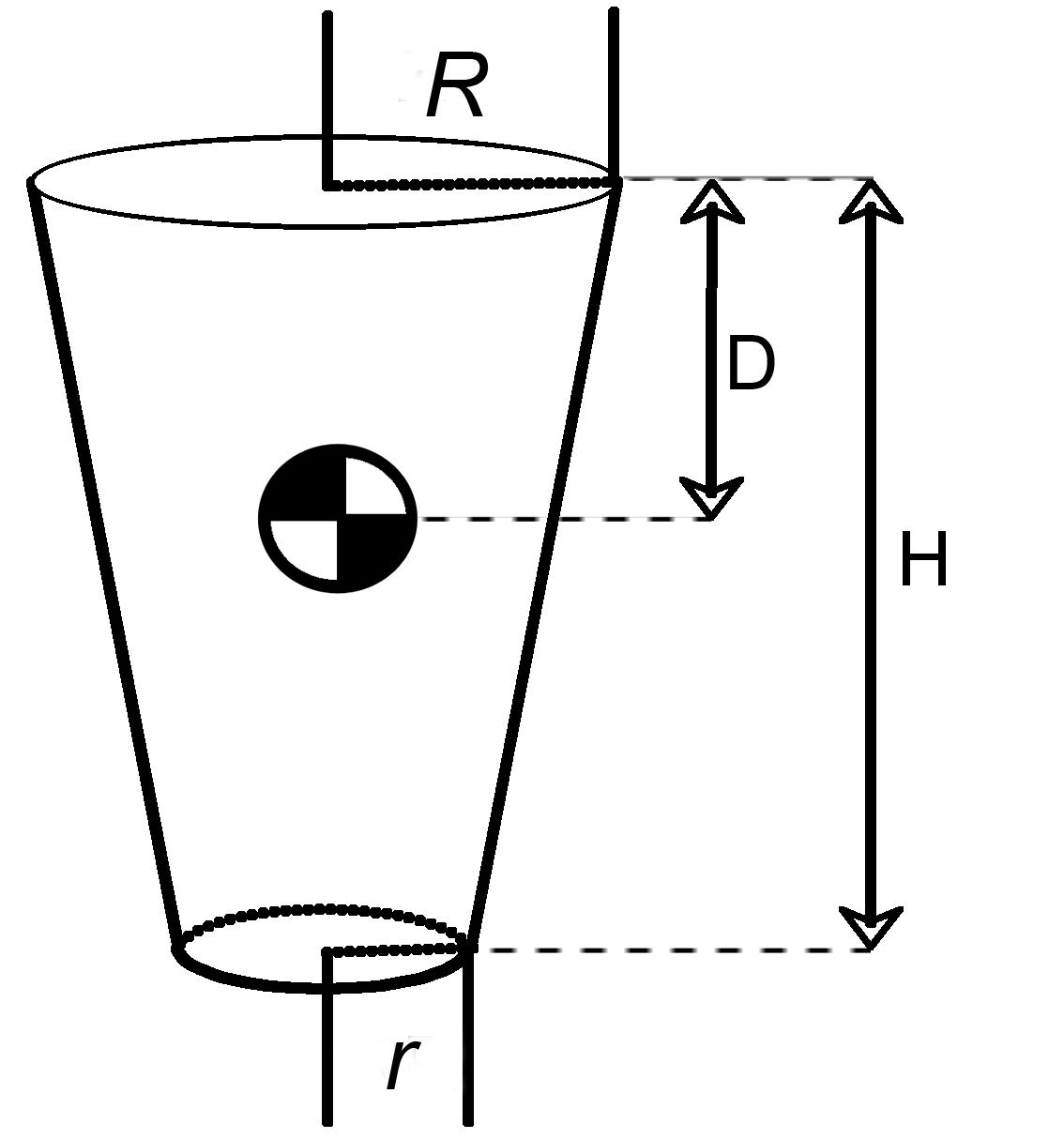

Considering:

, and	, then

and

where *ρ* is the volumetric mass set to 1000 kg/m^3^.

*r*

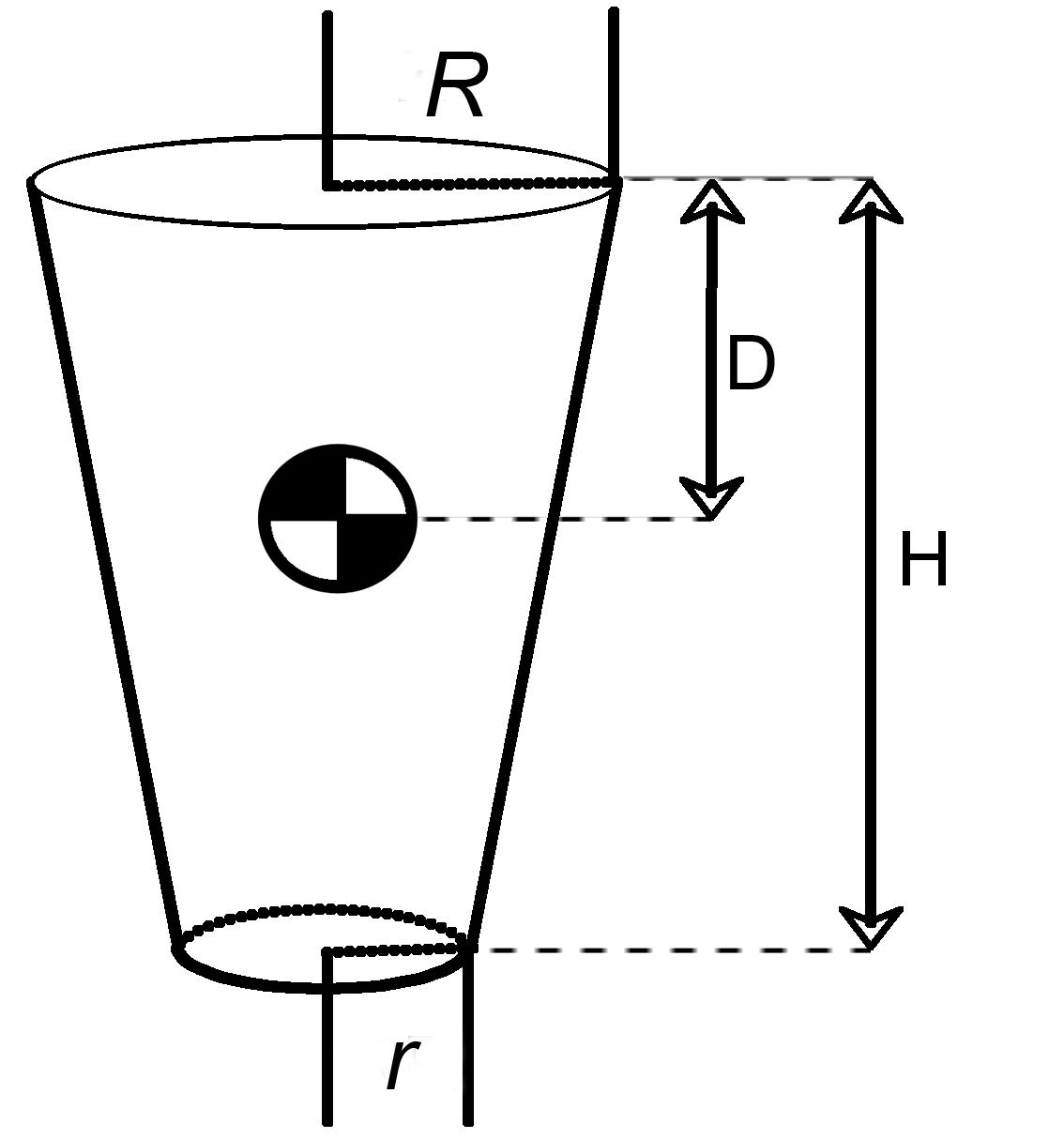

*L*

The trunk was considered as a cylinder, with the length *L* calculated from the average distance between the midpoint of the left and right shoulder and of the left and right hip (the midpoint was calculated as in Dewolf et al., (2018)). Knowing the mass (Table S1) and the volumetric mass, r can be computed as *r* =sqrt(V/(πL));

Then, I_0_ can be computed as: I=mass((r^2^/4)+(L^2^/12));

The head was considered as a sphere, with a diameter equal to 28% of *L* (this proportion was estimated on the photo of the elephants).

Table S1.

| Segment | *m*  (% of *M*_b_) | *R*  (m) | *r*  (m) | *H*  (m) | *D*  (%) | *I_0_*  (kg m^2^) |
| --- | --- | --- | --- | --- | --- | --- |
| Torso | 55.0 | - | - | - | - | - |
| Head | 16.0 | - | - | - | - | - |
| Upper Arm | 4.0 | - | 16.8 ± 1.9 | 66.9 ± 7.9 | 43.4 ± 0 | 4.75 ± 1.84 |
| Forearm | 2.0 | 16.8 ± 1.9 | 12.2 ± 1.2 | 63.5 ± 10.4 | 44.9 ± 1.9 | 1.82 ± 0.99 |
| Manus | 0.8 | 12.2 ± 1.2 | 17.2 ± 2.2 | 36.3 ± 3.9 | 55.4 ± 1.9 | 0.42 ± 0.15 |
| Total Fore | 6.8 | - | - | 166.8 ± 15.3 |  |  |
| Thigh | 5.0 | - | 18.4 ± 3.0 | 78.3 ± 8.8 | 43.4 ± 0 | 9.56 ± 5.40 |
| Shank | 2.0 | 18.4 ± 3.0 | 12.4 ± 1.6 | 67.6 ± 9.2 | 43.2 ± 1.6 | 2.41 ± 1.28 |
| Pes | 0.7 | 12.4 ± 1.6 | 15.9 ± 1.7 | 23.7 ± 2.0 | 54.3 ± 1.4 | 0.15 ± 0.05 |
| Total Hind | 7.7 | - | - | 169.6 ± 15.5 | - | - |

Where *R* and *r* respectively are the proximal and distal radius, *H* is the length of the segment. *D* is the distance between the proximal joint and the position of the centre of mass of the segment (in % of the segment length). *I_0_* is the moment of inertia of the segment, in kg m^2^. Values are average ± SD for 27 elephants.

**Sensitivity analysis.**

Variations in inertial parameter values (segment masses, centre of mass, and moments of inertia) can potentially affect the computation of Internal work. It has already been showed that errors in these estimates have only miniscule effects on inverse dynamics (Reinbolt et al., 2007; Ren and Hutchinson, 2008). Here, we evaluated how a ±15% changes of inertial parameters can change the results of internal work. We focused our comparison on the torso segment. As stated previously, the torso was considered as a cylinder, with the length *L* and a radius of *r.* Changing the measurement of trunk (either r and L) by +30% (comparing -15% and +15%) results in a 5±4% or 25±11% changes of W_int_ respectively. Changing the estimated volumic density by 30% results in a 11±8% change of W_int_. Changing I_0_ by 30% results in an increase of W_int_ on average by 6±14%.

**References**

**Dewolf, A. H., Ivanenko, Y., Zelik, K. E., Lacquaniti, F. and Willems, P. A.** (2018). Kinematic patterns while walking on a slope at different speeds. *J Appl Physiol (1985)* **125**, 642–653.

**Reinbolt, J. A., Haftka, R. T., Chmielewski, T. L. and Fregly, B. J.** (2007). Are patient-specific joint and inertial parameters necessary for accurate inverse dynamics analyses of gait? *IEEE Trans Biomed Eng* **54**, 782–793.

**Ren, L. and Hutchinson, J. R.** (2008). The three-dimensional locomotor dynamics of African (Loxodonta africana) and Asian (Elephas maximus) elephants reveal a smooth gait transition at moderate speed. *Journal of the Royal Society Interface* **5**, 195–211.
